# iProteinDB: an integrative database of *Drosophila* post-translational modifications

**DOI:** 10.1101/386268

**Authors:** Yanhui Hu, Richelle Sopko, Verena Chung, Romain A. Studer, Sean D. Landry, Daniel Liu, Leonard Rabinow, Florian Gnad, Pedro Beltrao, Norbert Perrimon

## Abstract

Post-translational modification (PTM) serves as a regulatory mechanism for protein function, influencing stability, protein interactions, activity and localization, and is critical in many signaling pathways. The best characterized PTM is phosphorylation, whereby a phosphate is added to an acceptor residue, commonly serine, threonine and tyrosine. As proteins are often phosphorylated at multiple sites, identifying those sites that are important for function is a challenging problem. Considering that many phosphorylation sites may be non-functional, prioritizing evolutionarily conserved phosphosites provides a general strategy to identify the putative functional sites with regards to regulation and function. To facilitate the identification of conserved phosphosites, we generated a large-scale phosphoproteomics dataset from *Drosophila* embryos collected from six closely-related species. We built iProteinDB (https://www.flyrnai.org/tools/iproteindb/), a resource integrating these data with other high-throughput PTM datasets, including vertebrates, and manually curated information for *Drosophila*. At iProteinDB, scientists can view the PTM landscape for any *Drosophila* protein and identify predicted functional phosphosites based on a comparative analysis of data from closely-related *Drosophila* species. Further, iProteinDB enables comparison of PTM data from *Drosophila* to that of orthologous proteins from other model organisms, including human, mouse, rat, *Xenopus laevis*, *Danio rerio,* and *Caenorhabditis elegans*.

## Introduction

Post-translational modification is essential for the regulation of many cellular processes. For example, phosphorylation can serve as a molecular switch for signal transduction (Beurel *et al.*, 2015; Hunter, 2000; Kockel *et al.*, 2010; Nagini *et al.*, 2018). Based on the annotation of PhosphoSitePlus (Hornbeck *et al.*, 2012; Hornbeck *et al.*, 2015), the average number of phosphosites per protein is twelve for the human proteome and seven for the mouse proteome. Evolutionary studies of protein phosphorylation have suggested that a significant fraction of these large numbers of phosphosites may be non-functional (Beltrao *et al.*, 2013; Landry *et al.*, 2009; Studer *et al.*, 2016) and that evolutionarily conserved phosphosites are often highly relevant for function (Studer *et al.*, 2016), as evidenced, for example, by the Mitogen Activated Protein Kinase (MAPK) or Extracellular Regulated Kinase (ERK) families (i.e. ERK/MAPK, JNK, p38). Generally the activation of these kinases requires phosphorylation within the sequence, TxY, residing within the “T loop” of the catalytic domain by an upstream MAPK-K/MEK kinase. Upon phosphorylation, the activation loop moves away from the active site, allowing substrate entry and phosphorylation. The TxY motif is conserved in the vast majority of MAPK/ERK family members, from yeast to man, allowing, for example, the generation of an antibody specific for the phosphorylated, active form of MAPK/ERK (Gabay *et al.*, 1997). Other examples of highly conserved phosphosites include ribosomal protein S6 (rpS6), which is conserved in essentially all organisms including yeast, plants, invertebrates, and vertebrates. The physiological roles of phosphorylation at Ser235/236 of rpS6 remained unclear until genetic approaches abolishing the phosphorylation sites were applied in model organisms (Meyuhas, 2015). These examples highlight how conservation can illuminate phosphosite function. Model organisms can play essential roles in the elucidation of the functions of post-translational modification of highly conserved sites.

Mass spectrometry (MS)-based proteomics is a powerful approach for large-scale identification and characterization of phosphorylation sites. Three large-scale *Drosophila melanogaster* phospho-proteomic datasets have been generated over the past years using MS. Two datasets were generated from cultured cells (Bodenmiller *et al.*, 2007; Hilger *et al.*, 2009) and one was generated from embryos (Zhai *et al.*, 2008). Because the coverage of each dataset is limited, and to further characterize the breadth of phosphorylation in *Drosophila*, we generated a new dataset for *Drosophila melanogaster* and five closely-related species: *D. simulans, D. yakuba, D. ananassae, D. pseudoobscura,* and *D. virilis.* To facilitate the use of this dataset, we built an online resource, iProteinDB, integrating our data with other large-scale PTM data (Bodenmiller *et al.*, 2007; Hilger *et al.*, 2009; Zhai *et al.*, 2008) and curated PTM annotations for *Drosophila* and other model organisms. At iProteinDB, users are able to align PTM data for any protein of interest from multiple resources, including data from the six *Drosophila* species, other model organisms, and human cells. Additional relevant information, such as disease-related protein variants, sub-cellular localization, and protein abundance during *Drosophila* development, is also provided at iProteinDB.

## Methods

### Generation of phosphoproteomics data

Pre-larval embryos of mixed sex and age from each of the six *Drosophila* species were collected. Since different species develop at different speeds, the timing of collection was different for each species. Flies were enticed to lay eggs by incubating in the dark on grape juice plates. Proteins from embryos lysed in 8 M urea were digested with trypsin and separated into 12 fractions by strong cation exchange chromatography. Phosphopeptides were purified with titanium dioxide microspheres and analyzed via LC-MS/MS on either an LTQ-Orbitrap or Orbitrap Fusion instrument (Thermo Scientific). SEQUEST was used for spectral matching. Peptides were filtered to a 1% FDR. Proteins were filtered to achieve a 2% final protein FDR (final peptide FDR near 0.15%) and a probability-based scoring method was used to assign the localizations of phosphorylation events (Beausoleil *et al.*, 2006). The reference genomes used for initial analysis are *D. mel* r5.53, *D. ana* r1.3, *D. pse* r3.1, *D. sim* r1.4, *D. vir* r1.2 and *D. yak* r1.3 from FlyBase. The sites were re-mapped to *D. mel* r6.16, *D. ana* r1.05, *D. pse* r3.04, *D. sim* r2.02, *D. vir* r1.06, *D. yak* r1.05 at iProteinDB.

### Predicting the probability of phosphorylation

We aligned the phosphorylation sites identified in our datasets from 6 *Drosophila* species based on orthologous relationships predicted by OMA (Altenhoff *et al.*, 2018; Altenhoff *et al.*, 2011; Altenhoff *et al.*, 2015). For each proteome, we assign the probability of a phosphoacceptor (S+T (together) and Y) to be phosporylated, using a two-step approach. First, we scan each proteomes to find the kinase specificity of each phosphoacceptor, using NetPhorest (Horn *et al.*, 2014). This provided 40 scores for kinases specificity for a given region. Then, a support vector machine algorithm (SVM-light) was trained on each of the six species, using all 40 scores. We extracted the surrounding region of sites that are detected to be phosphorylated, and they received an initial score of 1 (positive dataset). The regions surrounding non-detected phosphosites received an initial score of 0 (negative dataset). We sample the data (2000 for S+T and 800 for Y) to train the model, and then assign scores to unknown phosphosites based on the support vector machine output (detected phosphosites by MS always received a score 1, irrelevant of their prediction).

### Comparison of PTM data across major model organisms and human

Orthologous relationships of *Drosophila melanogaster* proteins to major model organisms including human, mouse, rat, *X. tropicalis*, zebrafish and *C. elegans* were obtained using DIOPT (release 7) with a DIOPT score of 3 or higher. Protein sequences of the best orthologous genes based on DIOPT score and each non-redundant isoform of *Drosophila melanogaster* gene were aligned using MAFFT (vs 7.305B). Observed PTM sites, conserved phosphorylation sites, domain and disease-related protein variants are annotated on the aligned protein sequences (Figure 1). To compare specific PTM sites, the sequence of a sliding window of five amino acids surrounding the identified phosphosite was extracted and compared across species. The number of identical amino acids was counted and the percent of identity was calculated by dividing the number of identical amino acids over the window length. Phylogenetic trees for protein kinases were generated with Jalview 2.10 (Waterhouse *et al.*, 2009) and illustrated in iTOL (Letunic and Bork, 2016).

**Figure 1.**
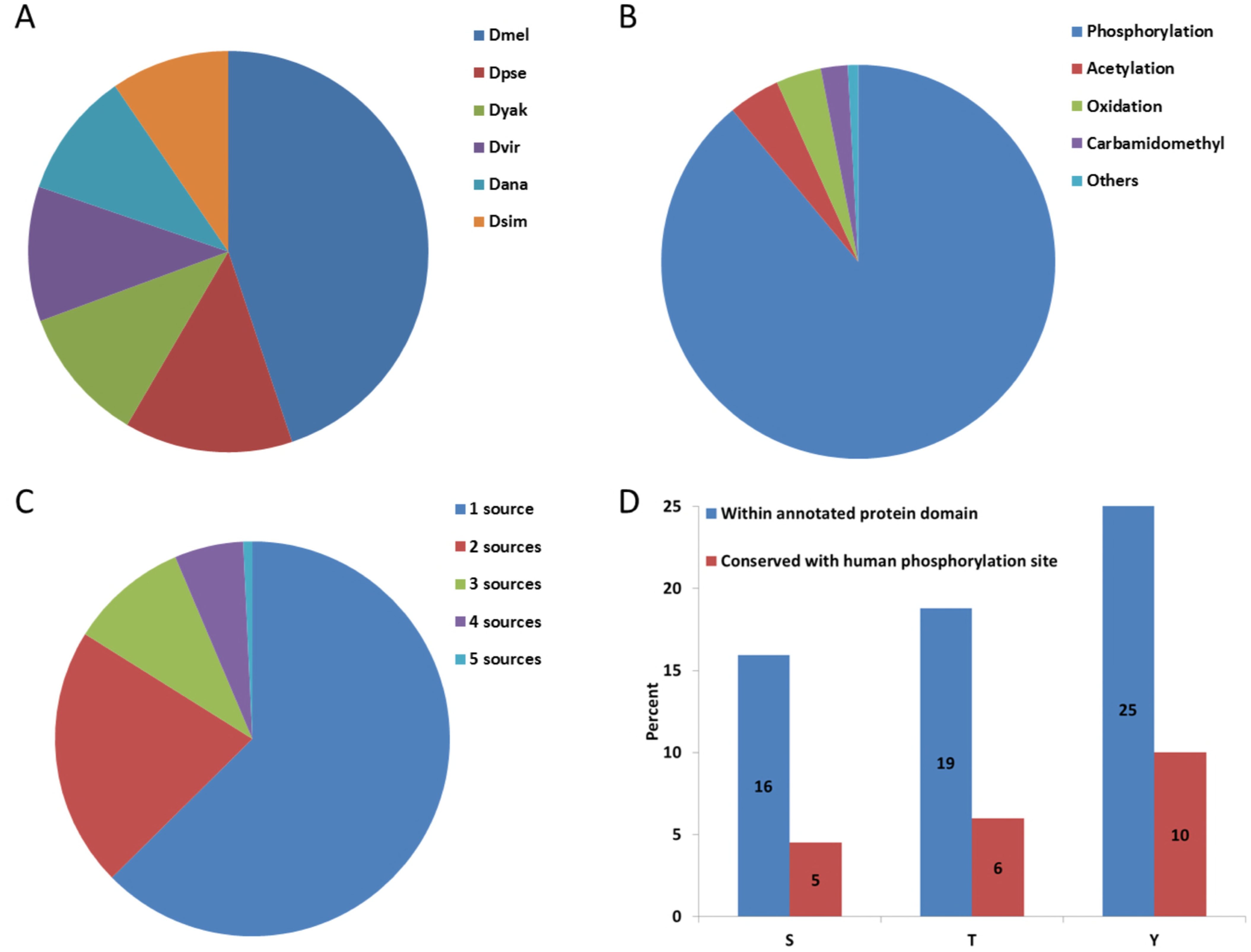
Database content and statistics. Distribution of 168,997 observed PTMs in the proteomics dataset (A). Representation of different types of PTMs (B). Overlap of phosphorylation data for *Drosophila melanogaster* from five different sources (C). Distribution of phosphorylation sites observed at three phospho-acceptor residues (serine (S), threonine (T) and tyrosine (Y)) within protein domains and their conservation based on at least 50% similarity to human sequence, considering a sliding window of five amino acids (D).

### Source of other data sets or tools

Protein information of 6 *Drosophila* species was obtained from FlyBase (ftp://ftp.flybase.net/releases/FB2017_03/). Protein information of human, mouse, rat, *X. tropicalis*, zebrafish and *C. elegans* were obtained from RefSeq (https://www.ncbi.nlm.nih.gov/refseq/). Other high throughput datasets of *Drosophila melanogaster* were obtained either from online resources (Phosida: http://141.61.102.18/phosida/index.aspx and Phosphopep: http://www.phosphopep.org/) or corresponding supplemental tables of relevant publications (Zhai *et al.*, 2008). The protein annotation of Swiss-prot and TrEMBL was downloaded from the UniProt FTP site (http://www.uniprot.org/downloads). Orthologous relationships were obtained from OMA (https://omabrowser.org/oma/home/) and DIOPT (http://www.flyrnai.org/diopt). Protein domain annotation of Conserved Domain Database (https://www.ncbi.nlm.nih.gov/Structure/cdd/cdd.shtml) was extracted from the RefSeq release files (.gbff files). Kinase motifs were predicted using the API of Scansite3 (http://scansite3.mit.edu/#home). PTM annotation of orthologous genes other than *Drosophila* was obtained from PhosphoSitePlus (https://www.phosphosite.org/staticDownloads.action).

### Implementation of the online resource

There were several steps involved that process the information and populate the back-end database of iProteinDB. After downloading data from relevant sources, such as UniProt, PhophoSitePlus and various publications, the extraction of relevant information was accomplished with in-house parsers written in Perl and Python. The redundancy of protein sequences was consolidated and a collection of distinct protein sequences from each *Drosophila* species was assembled based on FlyBase genome release (*D. mel* r6.16, *D. ana* r1.05, *D. pse* r3.01, *D. sim* r1.04, *D. vir* r1.02*, D. yak* r1.03). Since different resources annotate data based on different genome releases, we synchronized the data from various sources by mapping the original data (peptides) to the non-redundant protein collection of recent FlyBase genome release (see above) using the SeqIO interface of BioPython. Once filtered and updated, the data were then uploaded into a MySQL database, which is currently hosted by the Harvard Medical School (HMS) Research Computing group.

To display the data, we created a web-based application with PHP and a PHP framework called Symfony (version 2.6). Several client-side functions rely on JavaScript and AJAX, while some tabular displays use a jQuery plugin called DataTables.js, which allow for sorting and paging functionalities within the tables. This web application is also hosted by the HMS Research Computing group.

## Availability

iProteinDB is available for online use without any restrictions at https://www.flyrnai.org/tools/iproteindb/.

## Results

### Data integration and quality of six *Drosophila* phosphoproteomes

Embryos from six *Drosophila* species were collected, proteins were extracted and digested, phosphopeptides were isolated, and these samples were then subjected to ionization and fragmentation for identification and phosphosite determination using a mass-spec based method described previously (Sopko *et al.*, 2014). The data coverage ranges from 14,915 to 21,750 sites per species (Supplementary Table 1) and motif analysis of the data (Ullah *et al.*, 2016) shows that the most significant motifs of phosphosites in 6 *Drosophila* species are quite similar (Supplementary Figure 1). The orthologous relationships among the six *Drosophila* species, as well as other sequenced *Drosophila* species and mosquito species (Supplementary Figure 2), were predicted using the OMA algorithm (Altenhoff *et al.*, 2018), which infers orthologous genes among multiple genomes on the basis of protein sequence. Based on the multiple-sequence alignment of each orthologous group, the aligned positions were selected, for which phosphorylation was observed in at least one of the six *Drosophila* species. Given that the mass-spec based identification of phosphosites is incomplete, we filled the gap with machine learning predictions, using a similar approach as in (Studer *et al.*, 2016). A support vector machine (SVM) algorithm was trained to assign a propensity score of 0-1 to each corresponding phospho-acceptor residue (serine, threonine, or tyrosine) for each species for which that residue was not identified as phosphorylated, based on the likelihood of phosphorylation. This information is available at the iProteinDB resource (see below) to help researchers interested in identifying evolutionary conserved phosphorylation sites. We next compared the propensity score with the phosphoproteomics data from other sources. We found a strong correlation between the propensity score and the chance that a predicted site was phosphorylated as supported by independent datasets (Supplementary Figure 3a). We also compared the frequency of phosphorylation among the six *Drosophila* species with experimental data for orthologous human proteins. Not surprisingly, phosphorylation sites conserved among the six *Drosophila* species were more likely to be reported as phosphorylated at the corresponding sites in orthologous human proteins. This correlation was more prevalent for those sites with greater than 50% amino acid similarity between *Drosophila* and human orthologs (Supplementary Figure 3b).

We estimated the false negative rate for each of the six *Drosophila* species by selecting those sites that are 100% identical (considering eleven amino acid peptides comprising the phosphosite plus five amino acids upstream and downstream) among all six species and for which phosphorylation was observed in at least two species. The false negative rate is estimated to be the percent of the sites that are not covered by the data in each species. For example, 86% of these sites for *Drosophila melanogaster* are covered by at least one of the 4 datasets, and/or UniProt annotation so the false negative rate is about 14% while there is only 1 dataset for each of the other 5 *Drosophila* species, and therefore, the false negative rate is relatively higher, 44% to 79% (Supplementary Figure 3c).

### Integration of phosphoproteomes from other resources

We built the iProteinDB database to store phosphoproteomics data generated by our group and other large PTM datasets. Other datasets were obtained from the supplemental table of the original publications (Zhai *et al.*, 2008) or the relevant websites (Bodenmiller *et al.*, 2008; Bodenmiller *et al.*, 2007; Gnad *et al.*, 2011). Original data were mapped to the same version of the FlyBase proteome annotation (FB2017_03) and then integrated with our data in iProteinDB. The information of PTM sites and the score/peptide from the original source are stored and made available at the iProteinDB website. To compare PTM data across species, we integrated orthologous relationships of *Drosophila* species as predicted by OMA (Altenhoff *et al.*, 2018), the orthologous relationships among major model organisms predicted by DIOPT (Hu *et al.*, 2011), and PTM data for other species from PhosphoSitePlus (https://www.phosphosite.org) (Hornbeck *et al.*, 2012; Hornbeck *et al.*, 2015). The subcellular localization annotation and human disease related protein variants were integrated from UniProt (https://www.uniprot.org/), whereas protein domain annotation information was integrated from the National Center for Biomedical Information (NCBI) Conserved Domain Database (https://www.ncbi.nlm.nih.gov/cdd). Information about protein abundance during *Drosophila* development was also integrated from a recent publication (Casas-Vila *et al.*, 2017).

Altogether, iProteinDB covers 168,997 individual PTMs for *Drosophila*, of which 70,013 (41%) were observed in *Drosophila melanogaster* (Figure 1a). 62,239 (89%) of the *Drosophila melanogaster* PTM data collected in iProteinDB are phosphorylation sites, covering 8,068 unique proteins and 3,937 genes (Figure 1b). Comparing our *Drosophila melanogaster* phosphoproteomics data with that from other sources, we find that 61% of our data overlaps with one other source and 36% of our data overlaps with at least 2 other sources (Table 1).

Overall, 37% of the phosphorylation data is supported by multiple resources and thus can be considered high confidence (Figure 1c, Table 1).

## Online resource

Users can query *Drosophila* genes of interest, and choose one isoform if there are multiple non-redundant isoforms for the gene of the interest. There are three tabs from which to choose (Figure 2).

**Figure 2.**
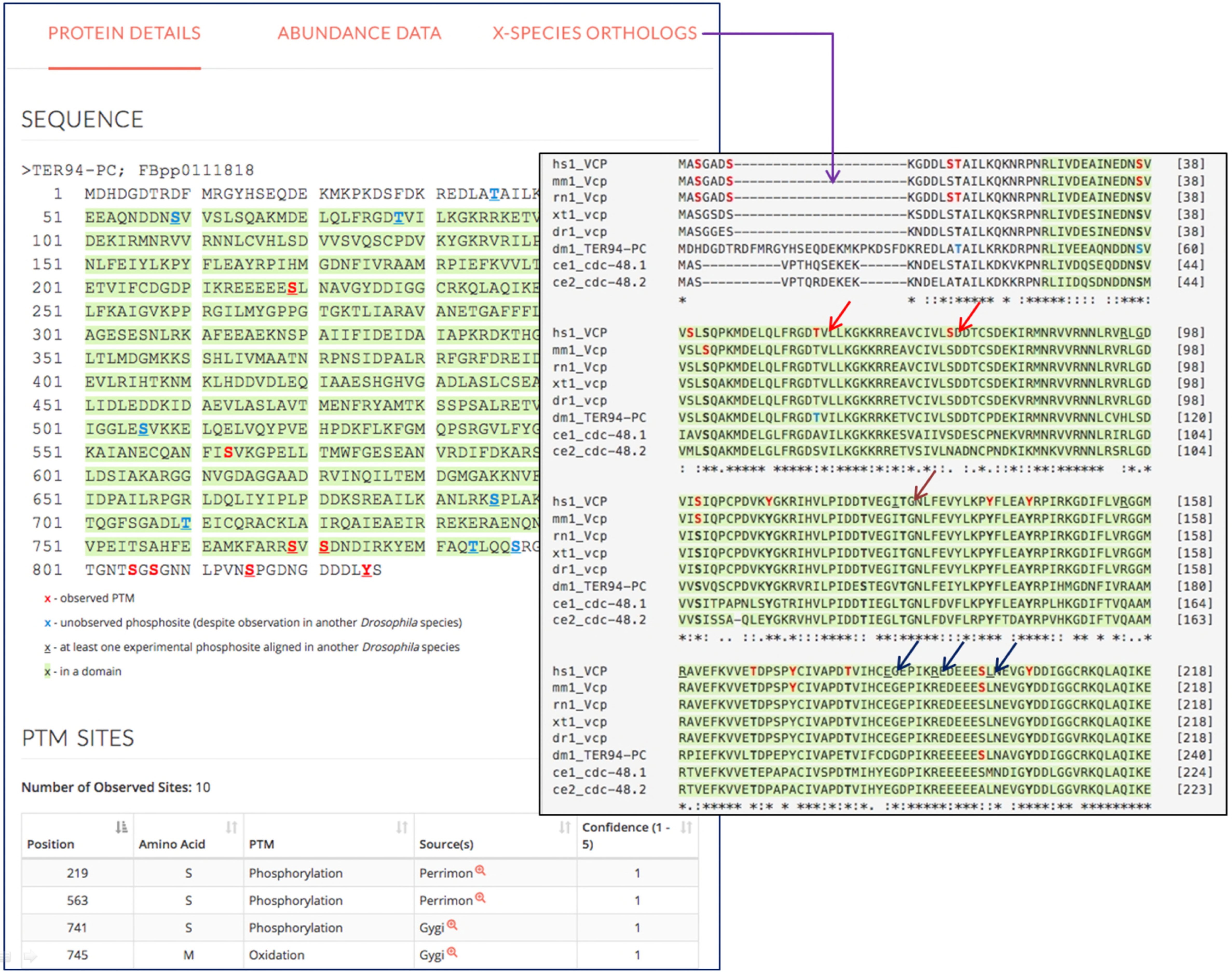
Features of iProteinDB user interface. Observed PTM sites are marked red on the *Drosophila melanogaster* protein sequence. Predicted phosphosites based on phospho-proteomic data from five other *Drosophila* species are marked in blue. Sites observed in more than one *Drosophila* species are underlined. The protein domains are highlighted in green. The data sources of PTMs are summarized. At the “Predicted Orthologs” page, the multiple sequence alignment of orthologous genes of major model organisms and human are displayed with observed sites color-coded (red arrows), conserved sites bolded (brown arrow) and human disease variant mutations underlined (navy arrows).

1.) Protein detail tab. A user can view the protein sequence from any of the 6 *Drosophila* species in FASTA format. PTM sites are color-coded. The amino acid is displayed in red if the PTM is observed or blue if it was not observed but is predicted to be phosphorylated based on the data from different *Drosophila* species. The amino acid is underlined if the phosphorylation event was observed in more than one *Drosophila* species. Protein domains are highlighted in green. A table summarizing all the PTM sites for a given protein, as well as the data sources from which the PTM information was extracted, is provided, along with detailed information from the original sources, i.e. the original scores and peptide sequences. A table summarizing all predicted sites based on data from closely related *Drosophila* species is provided with a link to detailed information and multiple sequence alignments. Also indicated in this tab is sub-cellular localization annotation from UniProt for each phosphoprotein and kinase predicted to act on individual sites, as identified using ScanSite3 (Obenauer *et al.*, 2003).

2.) Predicted ortholog tab. Users can find a table of the best ortholog candidates for major model organisms based on DIOPT ortholog predictions (Hu *et al.*, 2011). Multiple sequence alignments were performed based on the protein sequences of orthologous genes. The sequences of all the aligned *Drosophila* phosphosites, over a sliding window of five residues, were compared to the corresponding sequences of each orthologous gene and a similarity score was calculated by pair-wise comparison. For example, if 10 of the 11 amino acids (phosphorylation site plus five amino acids upstream and downstream) are identical between *Drosophila* and human sites, the similarity score was assigned as 0.9 (10 divided by 11). Then, an average similarity score was calculated based on all pairwise combinations at a given site. All phosphorylation sites with an average similarity score of >0.5 are listed and summarized as conserved sites. Human disease-related variants annotated at UniProt are also listed, along with sub-cellular localization annotation of all orthologous proteins from UniProt. Multiple sequence alignment (MSA) across major model organisms is displayed. For MSAs, observed PTM sites for all orthologous genes are color-coded, domains are highlighted, and disease variants are underlined. Conserved sites are bolded. As we hope that iProteinDB will lead to new discoveries and hypotheses on previously uncharacterized phosphorylation events, we further integrated information on availability of corresponding antibodies from Cell Signaling Technology for proteins and sites that are homologous between *Drosophila* and human to help users with experimental designs.

3.) Protein abundance tab. Protein expression levels from a comprehensive proteomic study covering the complete *Drosophila melanogaster* life cycle (Casas-Vila *et al.*, 2017) are plotted. On this tab, a user can view the stages of the *Drosophila* life cycle during which a protein of interest is expressed.

### The *Drosophila* kinomes and their substrates show significant evolutionary conservation

The integration of six *Drosophila* phosphoproteomes along with ortholog information enabled us to determine the conservation of the *Drosophila* kinome and to assess the evolutionary selective pressure on its substrates.

We found that the entire *Drosophila melanogaster* kinome as defined by Manning and colleagues(Manning *et al.*, 2002) shows orthologous counterparts in the other *Drosophila* species based on the OMA algorithm (Figure 3, Supplementary Figure 4). The few exceptions such as the absence of an orthologous Abl tyrosine kinase in *D. simulans* might trace back to poor genome sequence quality. Consistent with previous observations (Manning *et al.*, 2002) all *Drosophila* kinases showed strong evidence for orthologous counterparts in at least one of the six integrated model organisms. Only Tie-like receptor tyrosine kinase, Ack-like, and Wsck showed poor or no homology in eukaryotes other than *Drosophila* species based on DIOPT. The high conservation of the *Drosophila* kinome within flies and across other eukaryotes suggests that the corresponding substrates are also significantly conserved.

**Figure 3.**
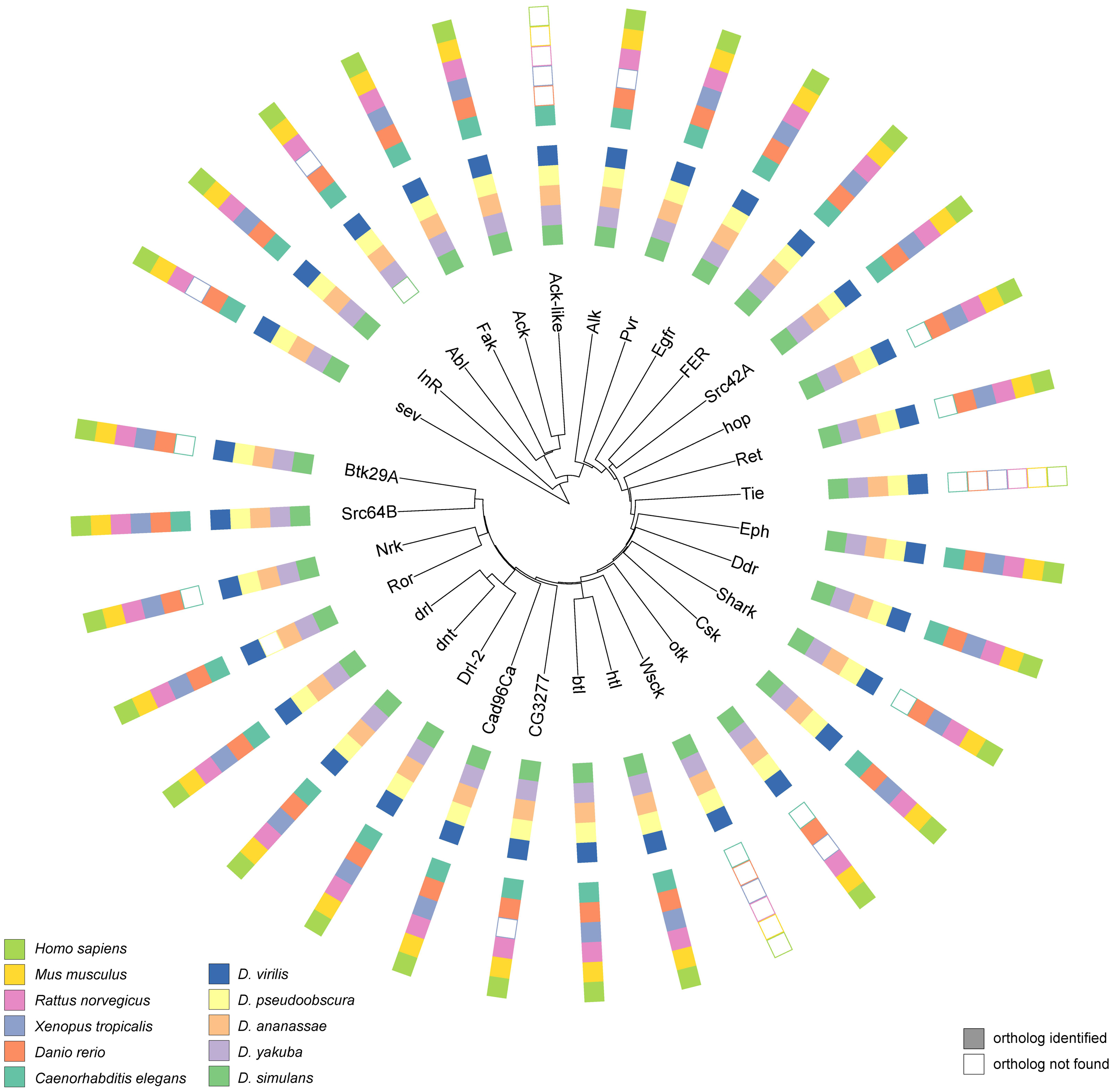
Evolutionary relationships among *Drosophila melanogaster* tyrosine kinases. The core of the plot illustrates the phylogenetic relationships between *Drosophila melanogaster* tyrosine kinases estimated by total sequence similarity. The outer circle reflects the presence of orthologs in other species.

To corroborate this hypothesis at the protein level, we determined the proportion of phosphorylated *Drosophila melanogaster* proteins that show homologies in other species. We used the conservation of *Drosophila melanogaster* proteins, which have not been found to be phosphorylated as a control to assess significance. We assume that this control set is indeed enriched for proteins that do not present kinase substrates. While phosphorylated and non-phosphorylated *Drosophila melanogaster* proteins have the same proportion of orthologs (99%) as the close relative *Drosophila simulan*, the phosphoproteome showed significantly higher conservation in more distantly related species from *Drosophila yakuba* (phospho: 99%, control: 97%; p < 0.01 based on two-sided Fisher Exact test) to *Caenorhabditis elegans* (phospho: 70%, control: 46%; p ^<^ 0.01) (Figure 4a). This suggests that not only the kinome but also its substrates are more conserved than other proteins. The significant conservation of the identified phosphoproteome might, however, also be partly driven by an enrichment of highly expressed proteins in the phosphoset and the presence of potential pseudogenes and predicted proteins in the control set. We therefore analyzed the conservation of phosphorylated versus non-phosphorylated residues of identified phosphoproteins, and found that phosphorylated residues show significantly higher conservation within all *Drosophila* species (p < 0.01) but not in more distant species (Figure 4b). Similar trends have been reported for other eukaryotic phosphoproteomes. For example, human phosphosites have been shown to have significantly higher conservation in mammals and other higher eukaryotes, but not in distant species including *Caenorhabditis elegans* or yeast (Gnad *et al.*, 2010). The prevalent localization of phosphorylation sites in fast evolving loop and hinge regions of proteins (Iakoucheva *et al.*, 2004) might make it difficult to map the associated site in aligned disordered regions of distantly related species. In contrast, non-phosphorylated serines, threonines, and tyrosines are not restricted to localization on the protein surface, and therefore tend to occur in more structured and slower evolving regions on the protein (Gnad *et al.*, 2007).

**Figure 4.**
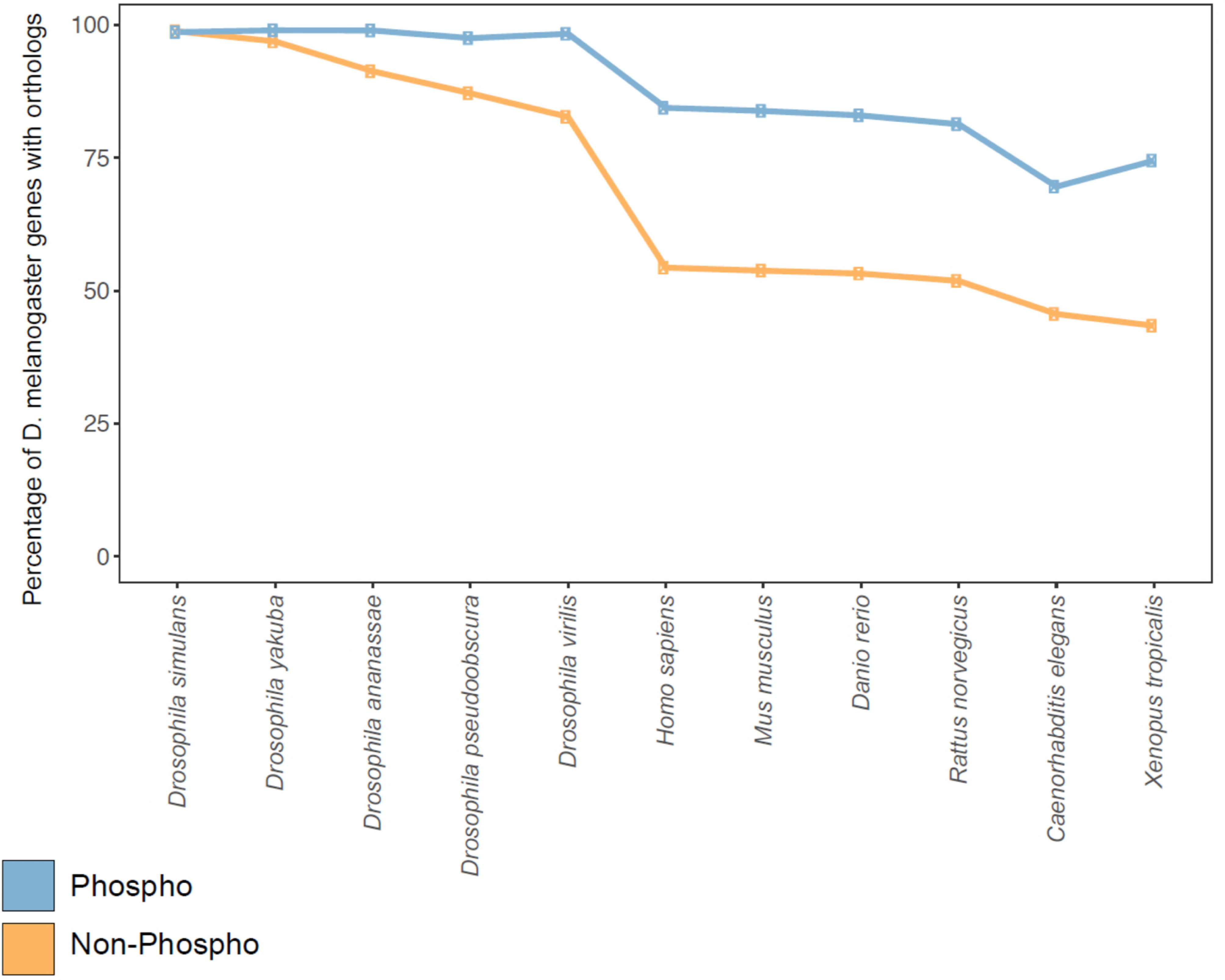
Conservation of phosphorylated proteins and sites. The line plot illustrates the proportions of *Drosophila melanogaster* phosphoproteins (blue) and non-phosphoproteins (orange) showing orthologs in other species (A). The line plot shows the proportions of conserved *Drosophila melanogaster* phosphosites (blue) and non-phosphorylated serines, threonines, and tyrosines (orange) across species (B).

In summary, we found significant conservation of the kinome and substrate proteins across all species. Similarly phosphorylated residues are significantly conserved within *Drosophila*, but difficult to trace back in distant species.

### Conservation between the *Drosophila* and the human phosphoproteomes underlines the utility of using the former as a model system

To assess the utility of the *Drosophila* phosphoproteome as a model for human phosphorylation events, we examined the evolutionary conservation of their phosphorylated sites with a focus on localization in functional domains and association with diseases. Approximately 17% of identified phosphosites are within annotated protein domains, and 82% of the identified phosphosites reside in proteins for which the corresponding *Drosophila* genes are conserved with human, based on DIOPT prediction using a score of 3 or more as cutoff (Hu *et al.*, 2011).

The corresponding human sites for 23% of these sites are also phospho-acceptors, among which, 3201 sites have 50% or more sequence identity to human sequence. Of the three phospho-acceptor residues, serine has the highest percentage of phosphorylation (Supplementary Figure 5) while phospho-tyrosine has the highest probability of residing within a defined protein domain and the highest sequence similarity with human orthologs (Figure 1d). Further analysis showed that the sequence identity of PTM sites between human and *Drosophila* melanogaster correlates with the probability that the associated phosphorylation event has also been observed in human cell phospho-proteomic datasets (Figure 5a).

**Figure 5.**
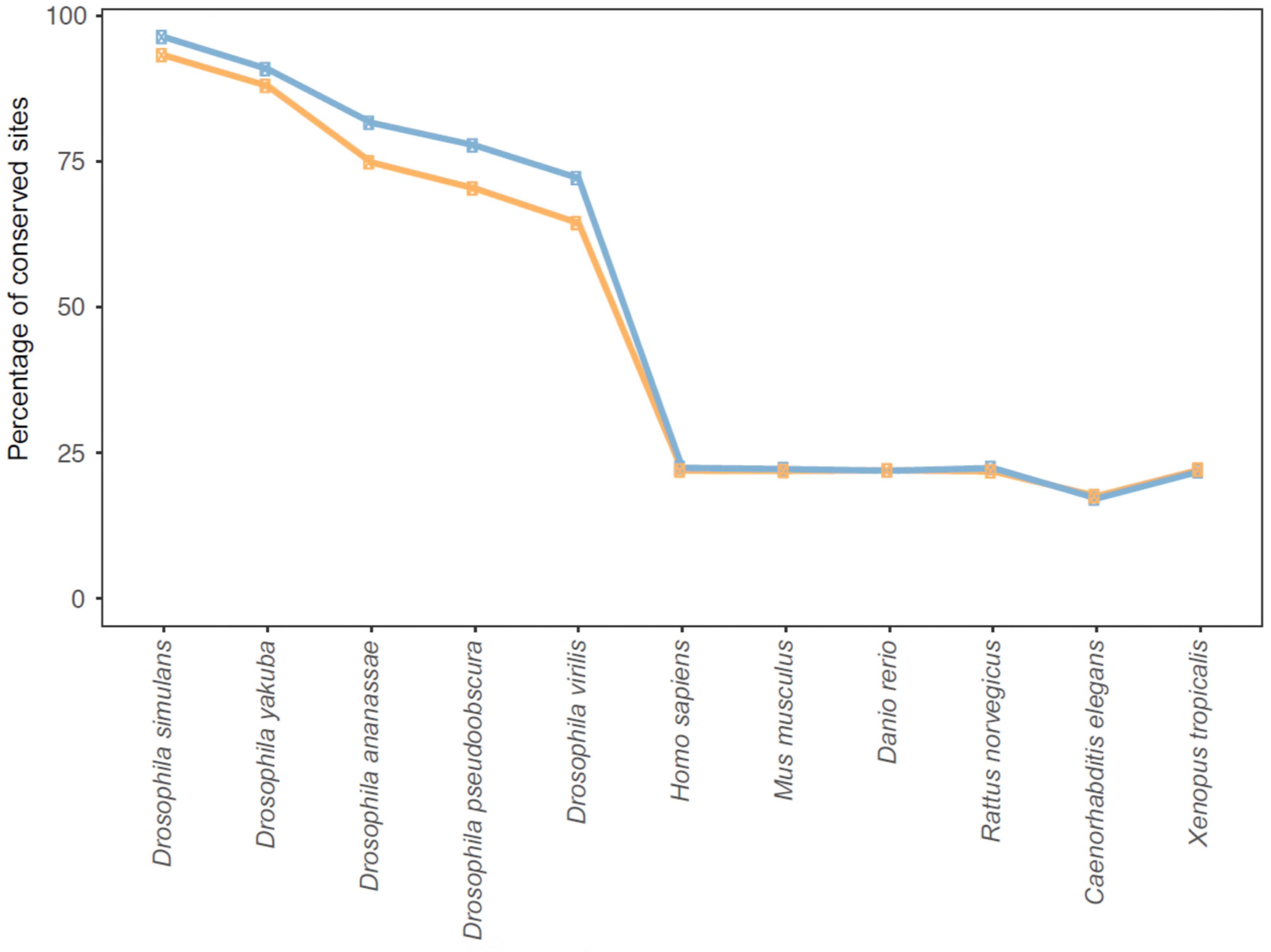
Analysis of the conservation of phosphorylation sites of *Drosophila melanogaster*. Correlations of sequence conservation and observed phosphorylation in *Drosophila melanogaster*: 11,619 phosphosites identified in *Drosophila melanogaster* proteins can be aligned to phospho-acceptor amino acids of the human orthologs, considering a sliding window of five amino acids surrounding the identified phosphosite. The probability of the corresponding phospho-acceptor site having been observed as phosphorylated in human data correlates with the degree of sequence similarity (A). Correlation of phosphorylation with disease related protein variants: The chance of the aligned human sites corresponding to the phosphosites identified in *Drosophila* locating within 10 amino acids distance to disease variants correlates with the sequence similarities between human and *Drosophila* sequences. The correlation is prevalent for phospho-acceptor sites in human (B).

We observed an enrichment of UniProt human disease related variants located proximal to phosphosites conserved with *Drosophila melanogaster*. For example, the enrichment p-value of disease related variants is 4.3*10^-10^ by Fisher exact test for the phosphosites with 80% or higher identity between human and *Drosophila* sites. To further analyze the intersection between phosphorylation events in *Drosophila* and disease variants in human, we calculated the percent of phosphosites that are within 10 amino acids of a disease variant at each identity cut-off. Our analysis indicates that more highly conserved sites tend to occupy positions proximal to residues variant in human disease (Figure 5b). For example, phosphosites with 50% or higher identity are about 2-fold more likely to be located within 10 amino acids of a disease variant than phosphosites with 20% or higher identity for the phospho-acceptor sites. Analysis of sites that are phospho-acceptor residues (serine, threonine, or tyrosine) in *Drosophila* but are not in human show a similar trend but this correlation was more prevalent for the phospho-acceptor sites, indicating that the correlation is driven by the phospho-acceptor as well as the conservation of the surrounding sequence.

Finally we compared all phosphosites in *Drosophila melanogaster* with their human orthologous sites. We identified 370 sites that were observed as phosphorylated in *Drosophila* and have 100% identity with human phosphosites over a sliding window of five amino acids. These sites cover 146 human genes, many of which are kinases, including cyclin dependent kinases, glycogen synthase kinases, mitogen-activated protein kinases, and ribosomal protein S6 kinases and the insulin receptor (InR) (Supplementary Table 2). For example, human glycogen synthase kinase 3A and 3B (GSK3A and GSK3B) auto-phosphorylate on a conserved tyrosine residue (Y279) for maximal activity, and play an important role in multiple signaling pathways (Beurel *et al.*, 2015; Nagini *et al.*, 2018). Dysregulation of GSK3 has been linked to various diseases including cancer, in which GSK3 can function as a tumor promoter or suppressor in different contexts and with different phosphorylation status (Ma, 2014; Nagini *et al.*, 2018; Sarkar *et al.*, 2015). The S278, Y279 and S282 sites within the protein-kinase domain of GSK3 have 100% identity with the *Drosophila* ortholog *sgg* (shaggy) and the phosphorylation of these sites has also been observed in *Drosophila* (Figure 6a). We further uncovered sites where the phospho-acceptor identity has changed, such as from serine to threonine (Figure 6b, Supplementary Table 3), suggesting that phosphorylation of these sites may also be conserved and required for regulation of the protein activity. We identified proteins for which the phospho-acceptor residues are conserved among *Drosophila* but absent in human despite the surrounding sequences being 100% identical (Figure 6c, Supplementary Table 3). These sites may regulate species-specific functions. Altogether these results indicate the utility of using *Drosophila* as a model system to study the function of these sites in signal transduction and the regulation of associated proteins.

**Figure 6.**
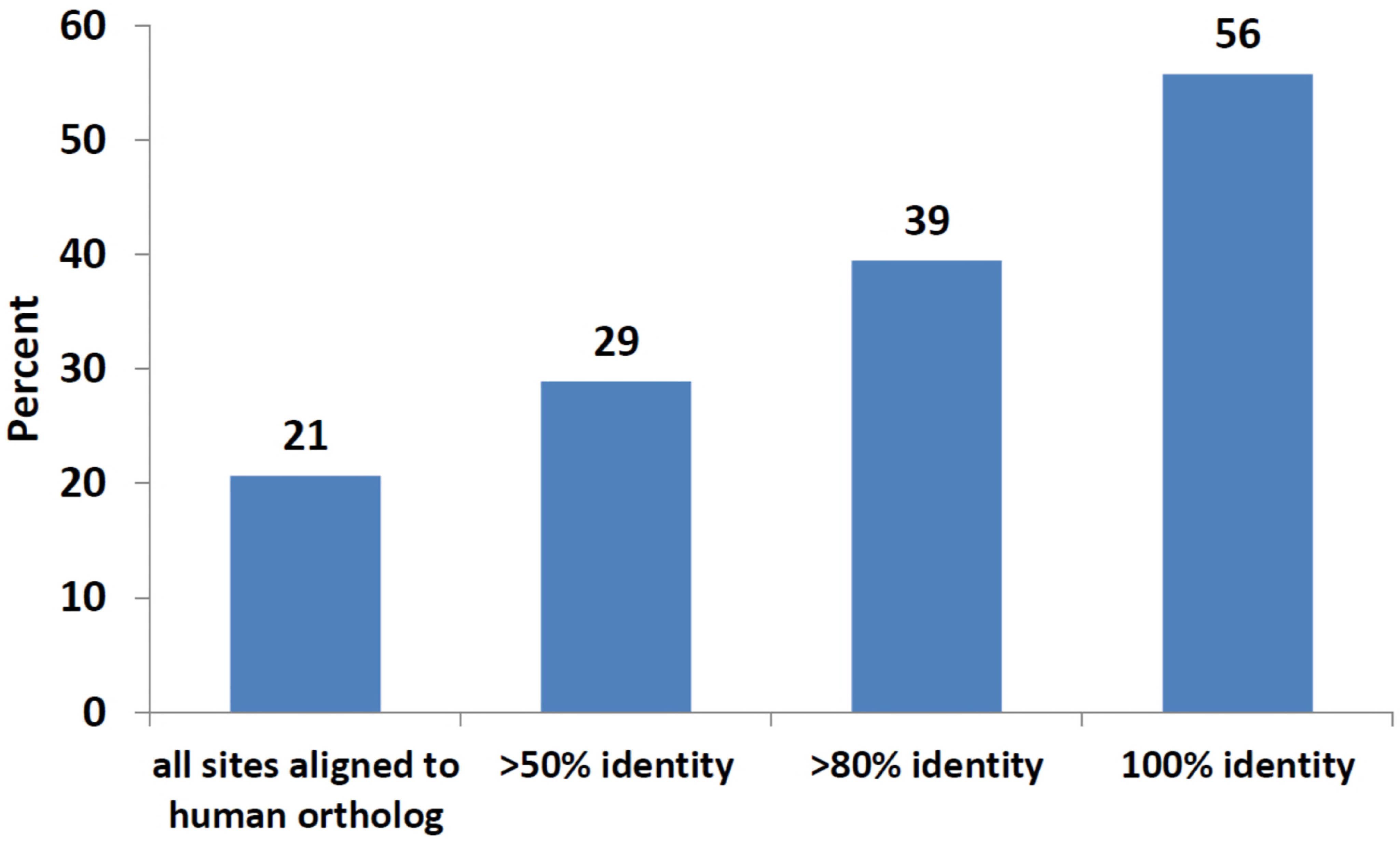
Examples of phosphosite conservation between human and *Drosophila melanogaster*. Examples of phosphosites identified in *Drosophila melanogaster* (red), also identified as phosphorylated in human (red), that share 100% identity with human (arrow) and indicated model organisms (A). Phosphosites where the observed phospho-acceptor residue has changed (B) and phosphosites where the phospho-acceptors have been lost but the surrounding sequences are 100% identical (C). The abbreviation of taxonomy name is used to represent different model organisms (hs - *Homo sapiens*; mm-*Mus musculus*; rn - *Rattus norvegicus*; xt - *Xenopus tropicalis*; dr-*Danio rerio*; dm-*Drosophila melanogaster*; ce-*Caenorhabditis elegans*).

## Conclusion

*Drosophila melanogaster* is one of the most-studied model organisms. Current PTM resources, such as PhosphoSitePlus (Hornbeck *et al.*, 2012; Hornbeck *et al.*, 2015), dbPTM (Huang *et al.*, 2016; Lee *et al.*, 2006) and Phospho.ELM (Diella *et al.*, 2004; Diella *et al.*, 2008; Dinkel *et al.*, 2011), have comprehensive coverage for human, mouse, and rat, but have very limited coverage for *Drosophila*. Resources like PHOSIDA (Gnad *et al.*, 2011), PHOSPHOPEP (Bodenmiller *et al.*, 2008; Bodenmiller *et al.*, 2007) and dbPAF (Ullah *et al.*, 2016) provide large-scale PTM data for *Drosophila* genes but are focused on only one or, at most, a few datasets. We generated a large-scale proteomics dataset of six closely related *Drosophila* species, made the data available, and integrated it with literature annotation and other large datasets for *Drosophila melanogaster*. This integrated resource allows researchers to obtain a more comprehensive view of the PTM landscape, taking into consideration all *Drosophila* proteomic data, and enabling comparison to orthologous proteins from other model organisms. Many of the conserved sites reside within kinases themselves, demonstrating that evolution has largely “optimized” protein kinase architecture and their operation within signaling pathways. We expect that iProteinDB will serve as a valuable resource to facilitate functional discovery. For example, iProteinDB can help a scientist identify sites that are critical for regulation that can be used for example to generate ‘activity-dead’ proteins that can serve as controls in rescue experiments with phosphomimetic (Pondugula *et al.*, 2009) and temperature-sensitive mutants (Hsu and Perrimon, 1994).

## Acknowledgments

We would like to thank the members of Perrimon laboratory, Gygi laboratory, *Drosophila* RNAi Screening Center (DRSC) and Transgenic RNAi Project (TRiP) for helpful input on the project. Particularly we would like to thank Dr. Stephanie Mohr for helpful suggestions during manuscript preparation and Mr. Aram Comjean for advice during the resource implementation. The DRSC is supported by National Institutes of Health (NIH) National Institute of General Medical Sciences grant R01 GM 067761 (to N.P.). The National Institutes of Health supported this work (5R01DK088718, 5P01CA120964, 5R01GM084947 and 5R01GM067761). R.S. is a Special Fellow of the Leukemia and Lymphoma Society. N.P. is a Howard Hughes Medical Institute investigator.

